# Nitric Oxide Modulates Auxin Signaling through TIR1 S-Nitrosylation During Thermomorphogenesis in Arabidopsis

**DOI:** 10.64898/2026.04.14.718228

**Authors:** NM Tebez, N Correa Aragunde, G Murcia, S Salvat, A Casco, DF Fiol, CA Casalongué, MJ Iglesias, MC Terrile

## Abstract

Auxin, a central hormone coordinating plant growth, integrates both environmental and developmental signals to regulate cell expansion, division, and organ patterning. Among these environmental cues, elevated ambient temperatures trigger a suite of developmental adaptations collectively known as thermomorphogenesis. In this study, we identify nitric oxide (NO) as a key mediator in the temperature-dependent regulation of auxin signaling. Our results show that warm temperatures (28–29 °C) enhance auxin-induced NO accumulation in *Arabidopsis thaliana* seedlings. Using pharmacological and genetic approaches, we demonstrate that NO is required for proper thermomorphogenic responses in aerial tissues. This redox signal promotes the stabilization and nuclear localization of the F-box auxin receptor TIR1, a crucial step for the activation of downstream auxin responses. Specifically, *tir1-1* seedlings expressing a non-nitrosylatable TIR1 variant mutated at the Cys140 residue exhibit impaired hypocotyl elongation and hyponasty under warm conditions compared to seedlings complemented with wild-type TIR1. These results highlight the functional relevance of the TIR1 Cys140 residue, a known target for S-nitrosylation, in coordinating thermomorphogenic responses. In contrast, the absence of TIR1 S-nitrosylation restricts primary root elongation at 22 °C but does not affect the growth-promoting effects of warm temperatures. Our findings uncover a novel redox-dependent regulatory layer in auxin signaling, where S-nitrosylation of TIR1 may modulate its stability and subcellular localization in a temperature- and organ-specific manner. This mechanism allows differential growth responses between shoot and root organs and highlights the complexity of hormonal and redox interplay during plant adaptation to elevated temperatures.

## 1. Introduction

Climate change projections predict a persistent rise in global temperatures, posing major challenges to plant growth, development, and productivity. Understanding how plants perceive and respond to elevated ambient temperatures is therefore crucial to anticipate and potentially mitigate the impacts of global warming on agriculture and ecosystems. One key adaptive strategy that plants use to cope with increased temperatures is thermomorphogenesis, which involves a suite of morphological and developmental responses that optimize plant architecture for thermal stress avoidance (Erwin et al., 1989). Within physiologically warm thermal ranges, *Arabidopsis thaliana* exhibits organ-specific growth changes. These include enhanced hypocotyl elongation (Gray et al., 1998), increased primary root growth (Hanzawa et al., 2013; Yang et al., 2017), and reduced cotyledon expansion. Additionally, warm temperatures induce hyponasty, an upward leaf movement driven by differential growth of the petiole between the abaxial and adaxial sides (Koini et al., 2009). This response is believed to improve leaf cooling by reducing light absorption (Crawford et al., 2012), thus contributing to plant thermotolerance.

Auxin, a central regulator of cell division and expansion (Perrot-Rechenmann, 2010), plays a pivotal role in thermomorphogenesis by integrating environmental and developmental signals. Elevated temperatures modulate multiple layers of auxin regulation, including biosynthesis, transport, perception, and signaling (Bianchimano et al., 2023). For instance, increased expression of *YUCCA* genes correlates with higher levels of free IAA in hypocotyls and root tips, facilitating growth in these organs. Warm environments also influence auxin homeostasis via epigenetic modifications and alternative splicing of transport-related genes. Canonical auxin perception relies on the TIR1/AFB family of F-box proteins, which act as auxin co-receptors and form part of the SCF^TIR1/AFB^ E3 ubiquitin ligase complex. In this signaling pathway, auxin functions as a molecular glue that promotes the interaction between TIR1/AFB proteins and the transcriptional repressors of the AUXIN/INDOLE-3-ACETIC ACID (Aux/IAA) family. Together, TIR1/AFB and Aux/IAA form the auxin co-receptor complex. Upon auxin binding, the SCF^TIR1/AFB^ complex mediates the ubiquitination and subsequent proteasomal degradation of Aux/IAAs, thereby releasing AUXIN RESPONSE FACTOR (ARF) transcription factors to activate or repress auxin-responsive genes. The proper function of this co-receptor system is essential for the temperature-induced promotion of hypocotyl and primary root elongation (Gray et al., 1998; Wang et al., 2016; Gaillochet et al., 2020), as well as for hyponastic leaf movement (van Zanten et al., 2009; Bianchimano et al., 2023). Interestingly, the expression of auxin receptors does not significantly change at the transcript level under warm conditions, suggesting that temperature influences their function via post-transcriptional or post-translational mechanisms (Bianchimano et al., 2023).

Previous studies demonstrated that warm temperature enhances TIR1 stability through the action of the molecular chaperone HSP90 and its co-chaperone SGT1 (Zhang et al., 2015; Wang et al., 2016). This chaperone-dependent mechanism ensures proper function of TIR1, thereby sustaining auxin responsiveness under warm conditions. Interestingly, nitric oxide (NO) signaling has also been implicated in temperature-responsive pathways that promote the accumulation of HSPs in plants, including HSP90 (Piterková et al., 2013; Manafi et al, 2021; Sanchez-Vicente et al., 2021). These observations suggest a potential connection between redox signaling and chaperone-mediated regulation of protein stability. Beyond chaperone-mediated stabilization, redox-based post-translational modifications have emerged as additional regulators of auxin perception. Among them, *S*-nitrosylation, a reversible covalent modification in which a NO group is attached to the thiol side chain of cysteine residues, represents a key mechanism through which NO acts as a signaling molecule in plants. NO is an important gasotransmitter that modulates diverse physiological processes. In *A. thaliana*, *S*-nitrosylation of TIR1 and its adaptor protein SKP1 enhances assembly of the SCF^TIR1/AFB^ complex and promotes receptor function (Terrile et al., 2012; Iglesias et al., 2018). Mutation of the conserved Cys140 residue in TIR1 disrupts its interaction with Aux/IAAs and compromises root development, highlighting the functional significance of this redox modification (Terrile et al., 2012). Recent evidence further indicates that redox modifications directly control TIR1/AFB2 nucleocytoplasmic dynamics. Lu et al. (2024) recently demonstrated in trichoblasts and root hair cells that auxin-triggered ROS and NO production drives oxidative modification (e.g., S-nitrosylation and other oxidative modifications) of TIR1/AFB2, promoting their nuclear localization and activation.

Despite these advances, the role of NO in modulating TIR1 function during thermomorphogenesis remains unclear. In this work, we demonstrate that warm temperatures enhance auxin-induced NO accumulation, thereby promoting TIR1 stabilization and nuclear localization at 29 °C. This redox-dependent regulation is essential for typical thermomorphogenic responses in shoots, such as hypocotyl elongation and hyponasty, while exerting no effect on primary root elongation. Moreover, we demonstrate that expression of a non-nitrosylatable TIR1 variant (tir1 C140A) fails to complement the *tir1-1* mutant in shoots fully, but retains functional activity in roots under warm conditions. Overall, our results uncover a novel, temperature-dependent layer of auxin signaling regulation via NO-mediated redox control of TIR1 stability and localization, positioning NO as a key modulator of shoot architecture during plant adaptation to warmer ambient temperatures.

## 2. Materials and Methods

### 2.1. Plant Material and Growth Conditions

*Arabidopsis thaliana* ecotype Columbia-0 (Col-0) was used in all experiments. The *tir1-1* mutant (Gray et al., 1998), *nia2* mutant (Wilkinson et al., 1991), *sav3* mutant (Tao et al., 2008), the reporter lines *DR5:GUS* and *IAA19:GUS* (Ulmasov et al 1997; Tatematsu et al., 2004), and the transgenic lines expressing TIR1-VENUS (Wang et al., 2016), TIR1-myc (Gray et al., 1999), or the non-nitrosylatable variant tir1-myc C140A in the *tir1-1* background (Terrile et al., 2012) were previously described. Seeds were surface-sterilized, stratified at 4 °C for 2-3 days, and germinated on vertically oriented Murashige and Skoog (MS) medium (0.5x MS salts, 1% sucrose, 0.8% agar, pH 5.8). Seedlings were grown under long-day conditions (16 h light/8 h dark, ∼100 μmol m^−2^ s^−1^) at 22 °C, or at 28-29 °C for warm treatment.

### 2.2. NO Donor and Scavenger Treatments

Endogenous NO levels were reduced using the scavenger cPTIO (2-phenyl-4,4,5,5-tetramethylimidazoline-1-oxyl 3-oxide), applied to the MS medium. Exogenous NO was supplied with S-nitrosocysteine (NO-Cys) as the NO donor, also added to the MS medium. Concentrations of cPTIO and NO-Cys varied depending on the assay and are indicated in each figure legend.

### 2.3. Hypocotyl, Root, Lateral Roots, and Hyponasty Measurements

For hypocotyl and primary root assays, seedlings were grown under long-day conditions at 22 °C for 3 and 4 days, respectively, before treatment. Seedlings were then subjected to the corresponding treatment for an additional 3d. Plates were photographed before and after treatment, and hypocotyl and primary root lengths were quantified with ImageJ software (ImageJ, version 1.53, NIH, Bethesda, MD, USA, https://imagej.nih.gov/ij/). For hyponasty assays, plants were grown for three weeks at 22 °C under short-day conditions to allow proper rosette development. Plants were then shifted from 22°C to 29°C for 4 hours, and images were taken before and after the temperature shift. The opening angle between the first pair of true leaves was measured using ImageJ/Fiji software. Each treatment included at least 15–20 seedlings or plants per biological replicate, and experiments were repeated at least three times independently.

### 2.4. GUS staining assays

For GUS expression analysis, seedlings were grown under specific conditions depending on the tissue or treatment analyzed: (i) auxin response in hypocotyls: 3-day-old IAA19:GUS seedlings were treated with 10 µM IAA and 1 mM cPTIO for 2 h; (ii) auxin response in roots: 4-day-old DR5:GUS seedlings were treated with 1 µM IAA and 1 mM cPTIO for 2 h; (iii) temperature and NO response: 3-day-old IAA19:GUS seedlings grown at 22 °C were transferred to either 22 °C (control) or 29 °C for 4 h, with or without 0.5 or 1 mM cPTIO. After treatment, seedlings were fixed in 90% acetone at −20 °C for 1 h, washed twice in 50 mM sodium phosphate buffer, pH 7, and incubated in staining buffer 50 mM sodium phosphate buffer, pH 7, 1 mg ml^-1^ 5-bromo-4-chloro-3-indolyl-b-D-glucuronide (X-Gluc, http://www.goldbio.com/), and stained for GUS activity from 2 h to overnight at 37 °C. Seedlings were cleared by an ethanol series (70, 50, 30, and 10%). Bright-field images were taken using a Nikon SMZ800 magnifier or Nikon Eclipse E200 microscope. GUS signal intensity was quantified using ImageJ software by measuring the mean gray value of the regions of interest, and results were expressed in arbitrary units.

### 2.5. Protein Extraction and Western Blotting

Total protein was extracted from 7-day-old seedlings using extraction buffer (50 mM Tris-HCl, pH 7.5, 150 mM NaCl, 10% glycerol, 0.1% Tween-20; 1 mM EDTA, protease inhibitor cocktail (Roche), and 50 μM MG132 to prevent proteasome-mediated degradation). Protein concentration was determined using the Bradford assay. Equal amounts of protein (30 μg) were separated by SDS-PAGE, blotted to nitrocellulose membranes, incubated with primary anti-GFP antibody (Agrisera, Sweden; 1:5000) overnight, followed by an HRP-conjugated secondary antibody (Invitrogen, United States) for 2 h. TIR1 was visualized using the enhanced chemiluminescence (ECL) kit (Amersham Biosciences). As a loading control, the large subunit of RUBISCO was visualized in membranes stained with Ponceau. Images were analysed using ImageJ software.

### 2.6. Confocal Microscopy

Confocal imaging was performed on a Zeiss LSM710 laser scanning microscope with a EC Plan-Neofluar 40x/1.3 oil immersion objective lens. For visualization of TIR1-VENUS, probes were excited with an Argon laser (λ = 514 nm), and fluorescence was collected at 524–602 nm. For chloroplast visualization, probes were excited with a He-Ne laser (λ = 633 nm), and fluorescence was collected at 643–735 nm. Transmitted light channel (T-PMT) was configured to visualize cellular structures. Hypocotyl images were processed with Fiji software. Fluorescence intensity was quantified from at least 10–15 nuclei and the surrounding cytosol per replicate, and the nucleus-to-cytosol fluorescence ratio was calculated. Briefly, the total fluorescence of the hypocotyl section was first measured, and the fluorescence corresponding to the nuclei within the same section was quantified separately. Cytosolic fluorescence was then estimated by subtracting the nuclear fluorescence from the total fluorescence of the section. The nucleus-to-cytosol fluorescence ratio was subsequently calculated using these values. The same quantification procedure was applied to all samples. Treatments (temperature shifts and/or cPTIO) are described in the corresponding figure legends.

### 2.7. NO Detection

Endogenous NO accumulation was assessed using the fluorescent probe DAF-FM DA (4-amino-5-methylamino-2’,7’-difluorofluorescein diacetate, Thermo Fisher). Four-day-old wild-type and *sav3* mutant seedlings were transferred to 22 °C or 29 °C for 2 h. DAF-FM DA (5 µM) prepared in 10 mM Tris-HCl buffer (pH 7.5) was added 30 min before the end of the temperature treatment, and seedlings were incubated in the dark. After incubation, samples were rinsed three times with the same buffer immediately before imaging. Fluorescence was visualized using an epifluorescence microscope with 488 nm excitation and 500–550 nm emission filters. Fluorescence intensity was quantified in whole hypocotyls using ImageJ/Fiji software.

### 2.8. Statistical Analysis

Data are presented as mean ± standard error (SE). Statistical analyses were performed using generalized linear models (GLMs). For most assays, a GLM with a Gamma distribution and log link function was applied, whereas for hyponasty measurements, data were transformed to values between 0 and 1 and analyzed using a Beta regression model. Post hoc comparisons were conducted with Tukey’s HSD test. Differences were considered significant at p < 0.05. All analyses and graphs were performed in R.

## 3. Results

### 3.1. NO acts as an intermediate in plant response to auxin in both the hypocotyl and root

Although auxin-induced NO production has been reported, and NO has been proposed to act as a second messenger in auxin signaling pathways in the roots of various plant species (Correa-Aragunde et al., 2004; Yadav et al., 2011; Terrile et al., 2012; Paris et al., 2018; Huang et al., 2022; Iglesias et al., 2024), the nature and extent of NO involvement in auxin-dependent processes remain to be fully elucidated. Therefore, we first examined the effects of scavenging endogenous NO using 2-(4-carboxyphenyl)-4,4,5,5-tetramethyl-imidazole-1-oxyl 3-oxide (cPTIO) on hypocotyl elongation and root development, as both are affected by elevated ambient temperatures. Exogenous auxin application enhanced hypocotyl elongation and increased lateral root density; however, both responses were significantly reduced in the presence of cPTIO (Fig. 1A-C). Supporting these findings, cPTIO also suppressed IAA-induced GUS expression in two auxin-responsive reporter lines: *pMASSUGU2 (MSG2)/IAA19:GUS,* which reports expression of the auxin-inducible gene IAA19 associated with hypocotyl elongation, and *DR5:GUS,* a synthetic auxin-responsive promoter widely used as a general marker of auxin signaling. This inhibitory effect was observed in both hypocotyl and root tissues, respectively (Fig. 1D-E, Supplementary Fig. S1; Pucciariello et al., 2018).

**Figure 1.**
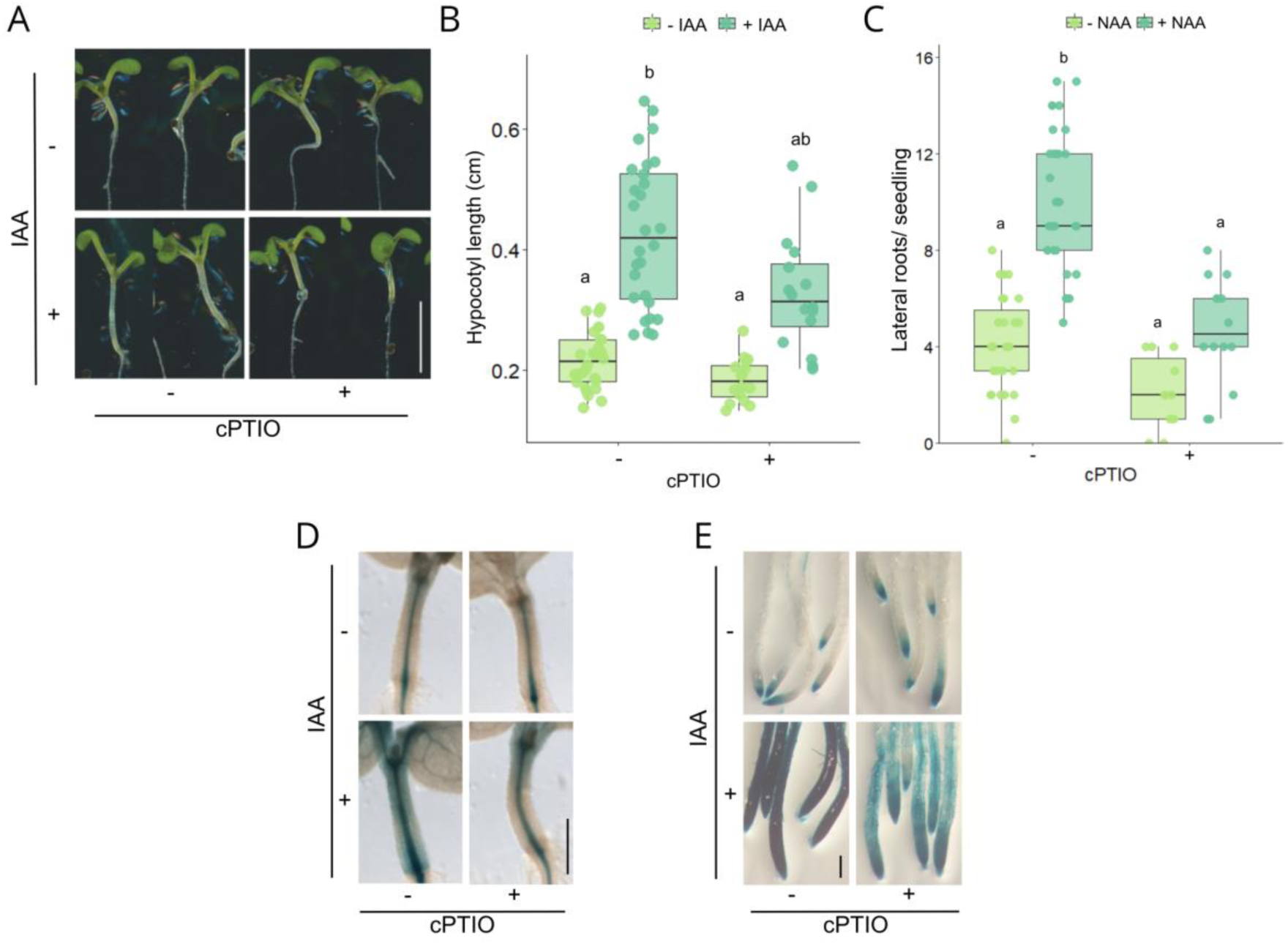
NO regulates auxin-related plant responses. **(A-B)** Three-day-old Col-0 seedlings were treated with IAA 5 μM, the NO scavenger cPTIO 0.5 mM, or both, and hypocotyl length was measured after 3 days. **(C)** Four-day-old Col-0 seedlings were treated with NAA 100 nM, cPTIO 1 mM, or both, and lateral roots were quantified after 4 days. (**D)** Three-day-old *IAA19:GUS* seedlings were treated with IAA 10 μM, cPTIO 1 mM, or both for 2 h and subjected to GUS staining.**(F)** Five-day-old *DR5:GUS* seedlings were treated with IAA 1 μM, cPTIO 1 mM, or both for 2 h and subjected to GUS staining. Boxplots in **(B)** and **(C)** represent the median and the first and third quartiles, and the whiskers extend to minimum and maximum values; all data points are shown as dots. Statistical differences were assessed using a generalized linear model (GLM, gamma distribution) followed by Tukey’s multiple comparison test (p < 0.05), as indicated by letters. Scale bars: 0.5 cm in **(A)**, and 0.2 cm in **(D, E)**.

### 3.2. NO is required for plant response to increasing mild temperatures

According to current models based on hypocotyl growth studies, warm temperatures promote auxin accumulation, which subsequently activates auxin-responsive genes and drives thermomorphogenic responses. Since endogenous NO is required for auxin-induced growth responses, we further investigated whether it also functions as a signaling molecule in thermomorphogenesis. We employed both genetic and pharmacological approaches to modulate NO levels. As expected, hypocotyl elongation was promoted at warm temperatures (29 °C) compared to ambient conditions (22 °C). However, this thermoresponsive growth was significantly reduced in *nia2*, a mutant in *NITRATE REDUCTASE 2,* which exhibits lower intracellular NO levels than wild-type plants (Fig. 2A), and also upon treatment with the NO scavenger cPTIO (Fig. 2B-C). To better understand the role of NO in auxin signaling at 29 °C, we examined the activation of auxin-responsive genes using the *MSG2/IAA19:GUS* reporter line. GUS expression was upregulated at 29 °C compared to 22 °C, with enhanced staining observed in hypocotyls (Fig. 2D, Supplementary Fig. S1). However, this induction was suppressed by cPTIO treatment, indicating that NO accumulation is required for proper auxin signaling activation under warm temperature conditions (Fig. 2D). Both hypocotyl elongation and auxin signaling activation were inhibited by cPTIO in a dose-dependent manner (Fig. 2B-D, Supplementary Fig. S2). In addition, the *nia2* mutant also exhibited a diminished leaf hyponastic response to warmth compared to control conditions (Fig. 2E-F), further supporting a role for NO as a novel regulatory signal in thermomorphogenesis, reinforcing the relevance of controlled intracellular NO level in mediating these temperature-responsive processes.

**Figure 2.**
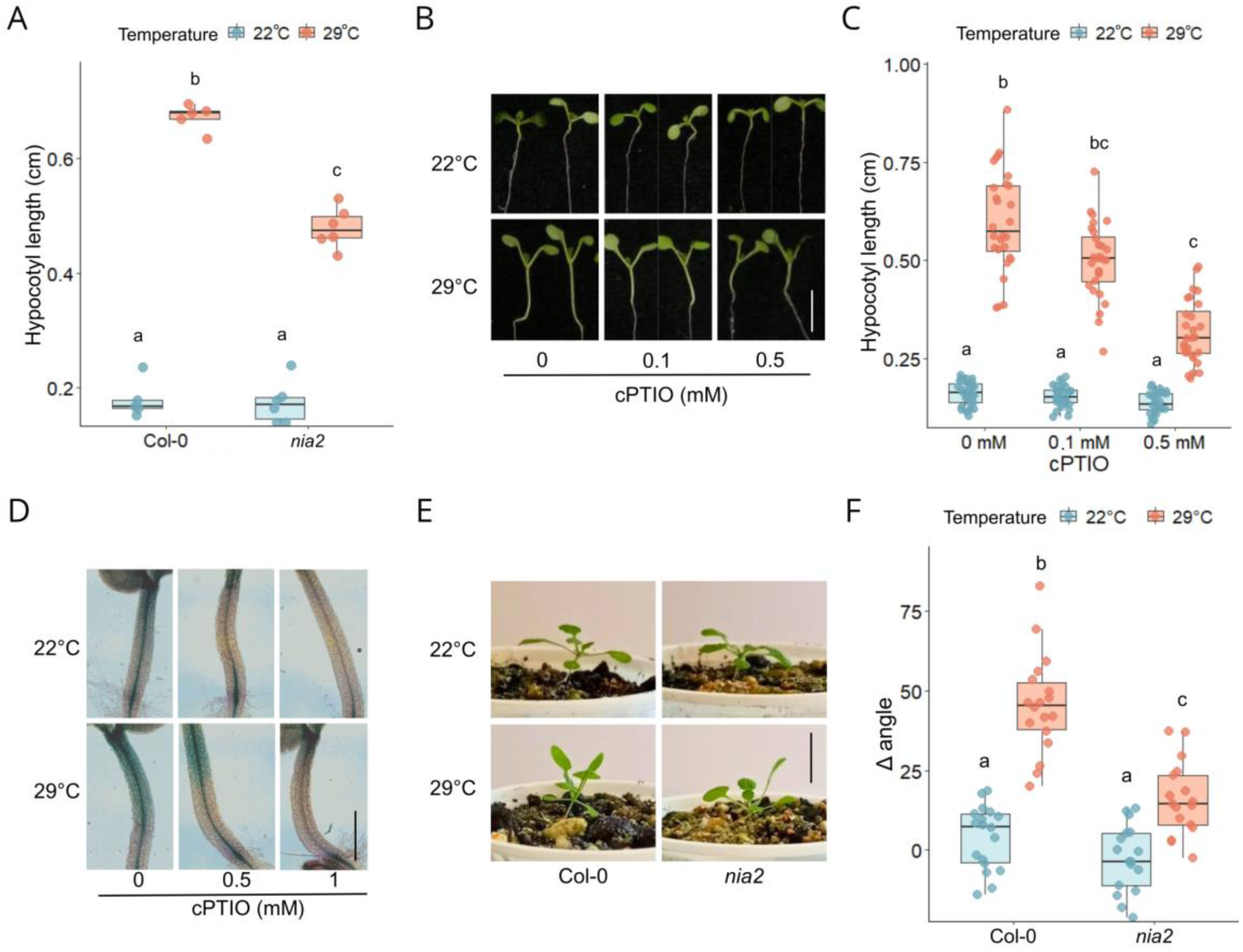
NO regulates auxin signaling during thermomorphogenesis. **(A)** Three-day-old Col-0 and *nia2* mutants were transferred to 22 °C or 29 °C, and hypocotyl length was measured after 3 days. **(B-C)** Three-day-old Col-0seedlings were treated with cPTIO and transferred to 22 °C or 29 °C; hypocotyl length was measured after 3 days. **(D)** Five-day-old *IAA19:GUS* seedlings were treated with cPTIO and transferred to 22 °C or 29 °C for 4 h and subjected to GUS staining. **(E-F)** Three-week-old Col-0 and *nia2* mutant plants were transferred to 22 °C or 29 °C for 4 h, and the opening angle of the first leaf pair of true leaves was measured before and after the temperature shift. Boxplots in **(A)**, **(C)**, and **(F)** represent the median and the first and third quartiles, and the whiskers extend to minimum and maximum values; all data points are shown as dots. Statistical differences were assessed using generalized linear models (GLMs): gamma distribution for panels **(A)** and **(C)**, and beta distribution for **(F)**. Tukey’s multiple comparison test (p < 0.05) was applied, and significant differences are indicated by letters. Scale bars: 0.5 cm in **(B)**, 0.2 cm in **(D)**, and 1 cm in **(E)**.

As a next logical step, we assessed whether seedlings increase NO levels when shifted from ambient to warmer temperatures. NO-associated fluorescence was monitored using the fluorescent probe DAF-FM DA, a widely used tool for detecting NO accumulation in plant tissues. Five-day-old seedlings grown at 22 °C were exposed to 29 °C for 2 h, and DAF-FM DA was added during the final 30 min of the treatment. Seedlings exposed to 29 °C displayed increased fluorescence in the hypocotyl compared with those maintained at 22 °C (Fig. 3A–B), indicating enhanced NO accumulation in response to warmth. Notably, this temperature-induced increase in fluorescence depended on auxin biosynthesis, as the *sav3* mutant (defective in the first step of the TAA/YUCCA auxin biosynthetic pathway) failed to show increased signal upon exposure to 29 °C (Fig. 3A-B). Consistently, *sav3* mutant seedlings show impaired hypocotyl elongation and hyponasty responses at 29 °C (Bianchimano et al., 2023). To explore potential molecular links between auxin signaling and NO production, and given that nitrate reductase (NR) activity has been implicated as a major source of NO downstream of auxin treatment (Kolbert et al., 2008ab; Cao et al., 2017), we examined the expression of the *NR* genes *NIA1* (AT1G77760) and *NIA2* (AT1G37130) across publicly available transcriptomic datasets (Supplementary Fig. S3). *NIA1* expression was consistently induced by auxin and also increased under elevated temperature conditions (27–28 °C) in seedlings. Although these data are correlative, they are consistent with a model in which auxin-dependent regulation of *NR* contributes to NO accumulation during thermomorphogenesis. However, the contribution of additional NO sources, including NOS-like activities, cannot be excluded.

**Figure 3.**
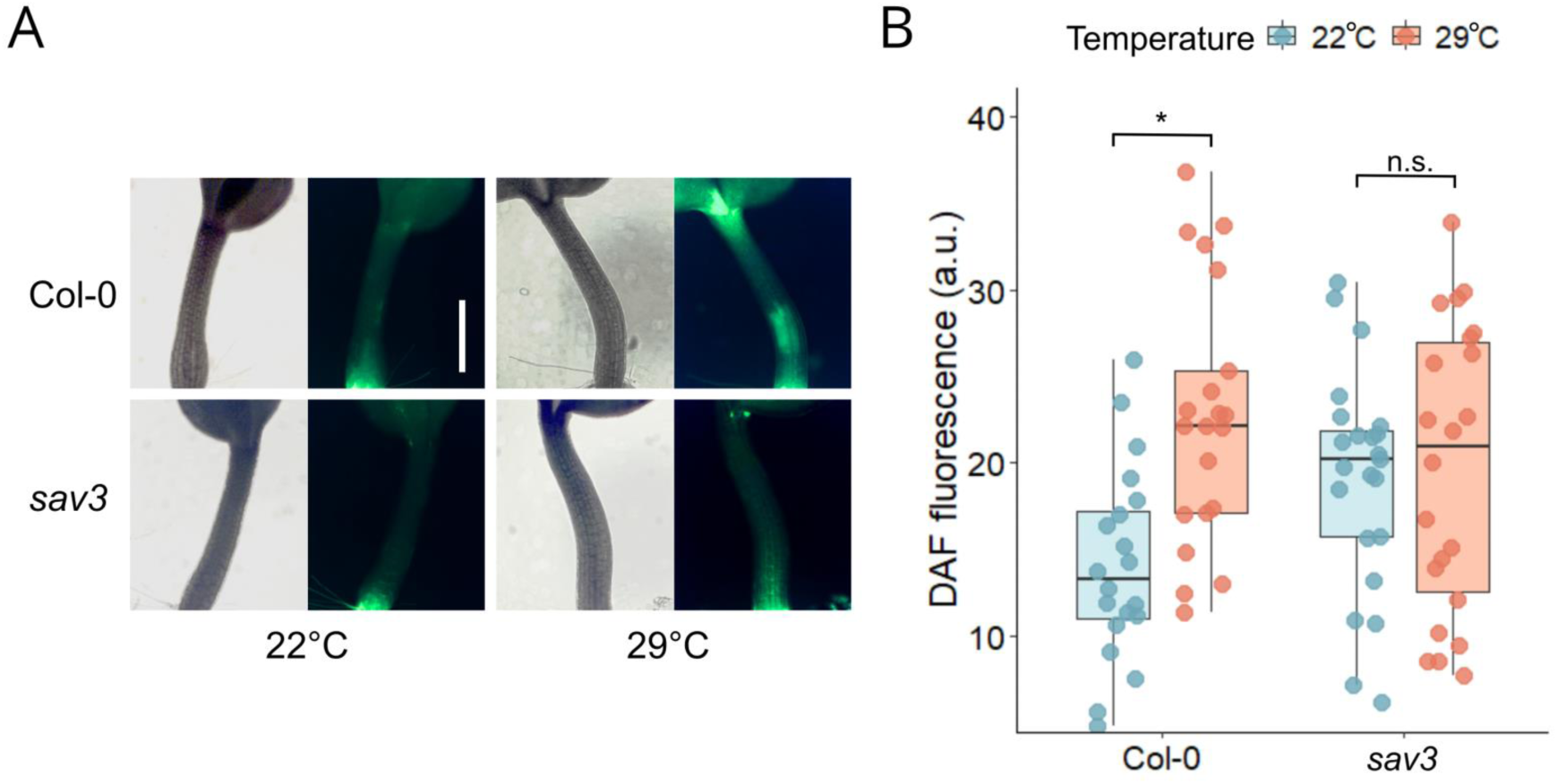
Temperature-dependent IAA biosynthesis induces NO accumulation. Four-day-old Col-0 and *sav3* mutant seedlings were transferred to 22 °C or 29 °C for 2 h. Subsequently, NO accumulation in the hypocotyls was quantified by DAF fluorescence. **(A)** Representative images of hypocotyls showing, for each condition, the bright-field image (left) and the corresponding DAF-FM DA green fluorescence (right) visualized using a GFP filter. Scale bar: 0.2 cm. **(B)** Boxplots represent the median and the first and third quartiles, and whiskers extend to the minimum and maximum values; all data points are shown as dots. Statistical differences were determined by pairwise t-tests (p < 0.05) and are indicated by different letters.

### 3.3 NO mediates nuclear TIR1 accumulation under warm temperatures

Previous studies have shown that TIR1 is rapidly stabilized in the nucleus at 29 °C, a process dependent on the HSP90 chaperone (Wang et al., 2016). Additionally, NO has been implicated in regulating TIR1 nuclear dynamics in root hair cells (Lu et al., 2024), as well as in the stabilization of TIR1 within the SCF E3 ubiquitin ligase complex (Iglesias et al., 2018). Based on this evidence, we investigated whether the nuclear accumulation of TIR1-VENUS in the hypocotyl of six-day-old *pTIR1:TIR1-VENUS* seedlings after a temperature shift to 29 °C depends on intracellular NO levels. As expected, we observed a substantial increase in nuclear TIR1-VENUS fluorescence upon exposure to 29 °C. However, this response was significantly attenuated by the NO scavenger cPTIO, indicating that NO accumulation is required for temperature-induced nuclear stabilization of TIR1 (Fig. 4A-B). Since previous studies reported that elevated temperatures do not affect the transcription of TIR1/AFB receptor genes (Wang et al., 2016; Bianchimano et al., 2023), we hypothesized that the observed increase in nuclear accumulation of TIR1-VENUS at 29 °C might be related to the regulation of protein stability. To test this, we examined whether NO and warm temperature treatments promote TIR1 protein stabilization by western blot analysis. TIR1 abundance was first assessed using an *in vitro* degradation assay with plant extracts incubated at 4 °C for 4 h in the presence of an NO donor (NO-Cys) and cPTIO scavenger (Iglesias et al., 2024). Treatment with NO-Cys prevented TIR1 degradation, whereas the addition of cPTIO counteracted the NO donor effect (Fig. 4C, E, and Supplementary Fig.S4). We included the proteasome inhibitor MG132 as a positive control, which also led to TIR1 stabilization. Analysis of *in vivo* TIR1 levels revealed a clear accumulation of the receptor at 29 °C compared to 22 °C control temperature (Fig. 4D, F). Additionally, treatment with the NO scavenger cPTIO under warm conditions led to a reduction in TIR1 protein abundance, suggesting that endogenous NO is required to maintain TIR1 stability (Fig. 4D, F). These findings further support the role of NO in regulating TIR1 stability and nuclear accumulation during thermomorphogenesis.

**Figure 4.**
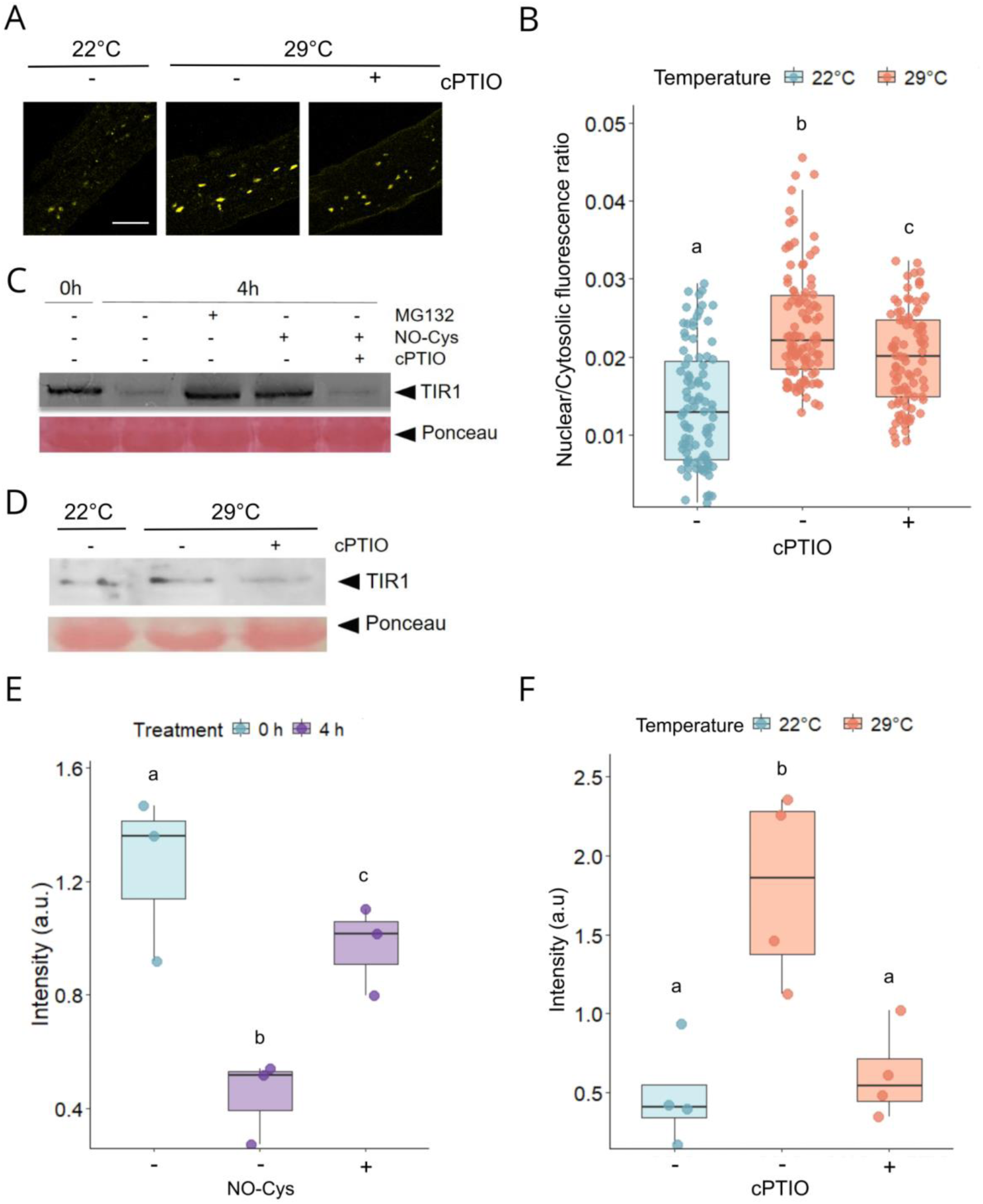
TIR1 nuclear accumulation at 29 °C depends on NO. Five-day-old *tir1-1 pTIR1:TIR1-VENUS* seedlings grown at 22 °C were treated with cPTIO 0.5 mM and subjected to 22 °C or 29 °C for 2 h. Hypocotyls were imaged using confocal microscopy **(A)**, and nuclear and cytosolic fluorescence intensities were quantified to calculate the fluorescence ratio **(B)**. Scale bar: 50 μm **(C, E)** *In vitro* degradation assays were performed using protein extracts from *tir1-1* 35S:TIR1-myc five-day-old seedlings incubated at 4 °C for 4 h in the presence of the NO donor NO-Cys, the NO scavenger cPTIO, or the proteasome inhibitor MG132. Protein extracts were analyzed by Western blot, and TIR1-VENUS levels were quantified relative to RUBISCO (Ponceau staining) used as a loading control. **(D, F)** Five-day-old pTIR1:TIR1-VENUS seedlings grown at 22 °C were treated with cPTIO 0.5 mM and then transferred to 22 °C or 29 °C for 2 h. Protein extracts were analyzed by Western blot, and TIR1 levels were quantified relative to RUBISCO, used as a loading control (Ponceau staining). Boxplots in **(B)**, **(E)**, and **(F)** represent the median and the first and third quartiles, and the whiskers extend to minimum and maximum values; all data points are shown as dots. Statistical difference according to ordinary one-way ANOVA coupled with Tukey’s multiple comparison tests (p < 0.05) is indicated by letters.

### 3.4. Non-nitrosylatable TIR1 mutant is impaired in the shoot thermomorphogenesis response

Under ambient temperature, NO exerts its regulatory role on TIR1, at least in part, through direct post-translational modification via S-nitrosylation at Cys 140 (Terrile et al., 2012; Terrile et al., 2022; Lu et al., 2024). To examine whether this regulatory residue is evolutionarily conserved, we performed a multiple sequence alignment of TIR1 homologs from diverse plant species, including several crop species. The cysteine corresponding to Arabidopsis TIR1 Cys140 was found to be highly conserved across these lineages, including several species of agronomic relevance (Supplementary Fig. S5). This conservation suggests that redox regulation of auxin perception may represent a broadly conserved feature of TIR1 function in plants.

To assess the functional requirement of the Cys140 residue in thermomorphogenic responses, we examined a range of growth phenotypes associated with warmth in a non-nitrosylatable mutant. Under warm conditions (29 °C), *tir1-1* mutants exhibited significantly reduced hypocotyl elongation compared to wild-type seedlings (Fig. 5A; Wang et al., 2016). This phenotype was fully complemented by the introduction of the wild-type *TIR1* gene, confirming the role of TIR1 in mediating hypocotyl growth at 29 °C. In contrast, the tir1 C140A variant, which carries a point mutation that replaces the nitrosylation target cysteine with alanine, was unable to restore the warm-temperature growth response, indicating that the S-nitrosylation site at Cys140 is essential for TIR1 function during hypocotyl elongation under warmth (Fig. 5A). Furthermore, to evaluate the impact of TIR1 S-nitrosylation beyond hypocotyl elongation, we analyzed temperature induced leaf hyponasty. Plants expressing the non-nitrosylatable tir1 C140A variant displayed a markedly reduced hyponastic response at 29 °C compared to those expressing the wild-type TIR1 protein (Fig. 5B). These results collectively support the hypothesis that the Cys140-dependent regulation of TIR1 is a critical mechanism through which NO modulates shoot-related thermomorphogenic responses.

**Figure 5.**
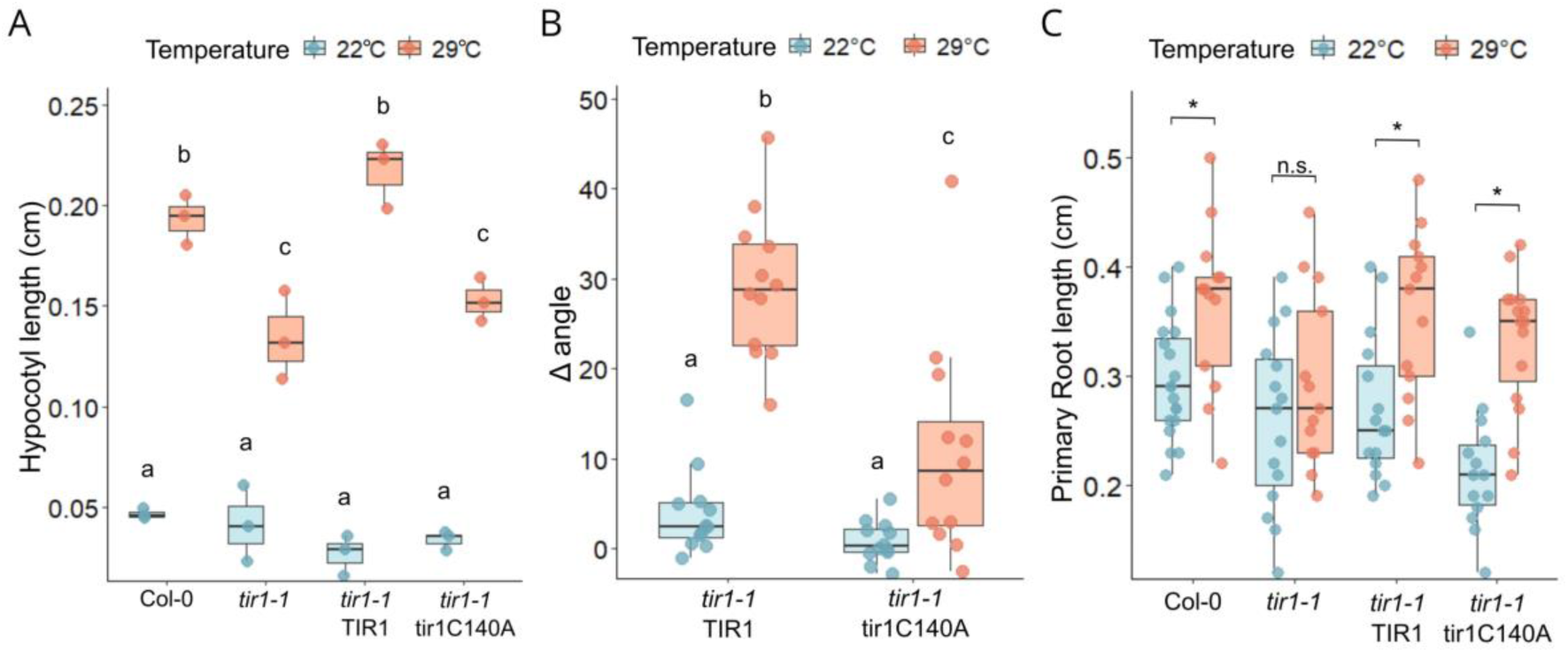
Non-nitrosylatable tir1 mutant is impaired in shoot thermomorphogenesis responses. **(A)** Four-day-old seedlings were transferred to 22 °C or 29 °C, and hypocotyl length was measured after 3 days. **(B)** Three-week-old plants grown at 22 °C were shifted to 22 °C or 29 °C for 4 h. The opening angle of the first leaf pair was measured before and after the temperature shift. **(C)** Four-day-old *tir1-1* TIR1 and *tir1-1* tir1 C140A seedlings were transferred to 22°C or 29°C, and primary root length was measured after 2 days. Boxplots represent the median and the first and third quartiles, and the whiskers extend to minimum and maximum values; all data points are shown as dots. Statistical differences were assessed using a generalized linear model (GLM, gamma distribution) followed by Tukey’s multiple comparison test (p < 0.05), as indicated by letters.

### 3.5. TIR1 S-nitrosylation is not required for primary root elongation at 29 °C

Warm temperatures promote auxin signaling activation through TIR1 perception which impact on primary root elongation (Wang et al., 2016); accordingly, the *tir1-1* mutant displays reduced growth at 29 °C compared to wild-type seedlings (Fig. 5C). To assess the functional relevance of TIR1 S-nitrosylation in the thermomorphogenic response at the root level, we tested whether the *tir1-1* phenotype could be complemented by either wild-type TIR1 or a non-nitrosylatable tir1 C140A variant. As shown in Fig. 5C, the tir1 C140A protein was able to complement the *tir1-1* mutant and restore primary root elongation at warmth. Interestingly, while the Cys140 mutation appeared to impair root growth at 22 °C, this effect was mitigated at 29 °C, suggesting that additional regulatory mechanisms may compensate for the lack of S-nitrosylation under warm conditions.

## 4. Discussion

Plant adaptation to elevated ambient temperatures involves complex signaling networks that integrate hormonal pathways with environmental and cellular cues. Auxin plays a central role in orchestrating thermomorphogenic growth responses, and our study identifies NO as a key redox signal that modulates auxin perception by promoting the stability and nuclear accumulation of the auxin receptor TIR1 under warm conditions. Our results show that warmth stimulates auxin-dependent NO production (Fig. 3), which correlates with increased nuclear accumulation and stabilization of TIR1 (Fig. 4). This promotes auxin-dependent elongation of the hypocotyl and the abaxial side of the petiole during thermomorphogenetic responses in aerial tissues (Fig. 2). Previous studies have shown that S-nitrosylation of TIR1 in Cys140 residue enhance auxin signaling by promoting SCF^TIR1–AFB^ complex assembly, stabilizing the receptor, and favoring its nuclear localization (Terrile et al., 2012; Iglesias et al., 2018; Lu et al., 2024; Yu et al., 2015). Consistent with a redox-dependent mechanism, seedlings expressing the non-nitrosylatable tir1C140A variant exhibit impaired shoot thermomorphogenic responses (Fig. 5). In this context, the phenotypes observed in plants expressing the non-nitrosylatable tir1C140A variant provide genetic evidence supporting a role for redox regulation of TIR1 in thermomorphogenic growth. Together, these observations uncover a previously unrecognized post-translational regulatory layer within the auxin signaling pathway, identifying NO as an important modulator that links temperature perception with auxin receptor function during thermomorphogenic growth.

Our data support a model in which NO intersects with chaperone-dependent mechanisms that regulate auxin receptor stability. Consistent with this, warmth-induced stabilization of TIR1 depends on HSP90 and its co-chaperone SGT1 (Wang et al., 2016; Zhang et al., 2015), and our results further indicate that NO contributes to TIR1 stabilization, suggesting convergence between redox signaling and chaperone-dependent pathways at the receptor level. To explore whether structural features of TIR1 could facilitate such cooperation, we analyzed AlphaFold3-based predictions of the TIR1–HSP90.2 interaction, including explicit modeling of S-nitrosylation at Cys140 (Supplementary Fig. S6). Comparison of modified and unmodified models revealed localized differences at the predicted interaction interface, including a transition of a region in HSP90.2 from a flexible loop to a more ordered α-helical conformation in the presence of the modification, together with a modest increase in interface confidence (ipTM). While these structural predictions do not establish a mechanistic effect, they suggest that redox modulation at Cys140 may favor a conformational state of TIR1 that is more compatible with HSP90 engagement, potentially enhancing chaperone–client stability. This interpretation is further supported by the proximity of the *tir1-1* mutation (G147D) to the Cys140 site and by previous observations that HSP90.2 overexpression can compensate for this defect (Watanabe et al., 2016). Together, these findings are consistent with a scenario in which redox-dependent and chaperone-mediated pathways converge on a shared structural region to regulate TIR1 stability during thermomorphogenesis, although experimental validation will be required to establish the mechanistic basis of this interaction.

Beyond its role as a redox-sensitive regulatory node, recent work has expanded the signaling repertoire of auxin receptors by revealing that TIR1 and other AFB proteins also possess auxin-activated adenylate cyclase activity, generating cAMP as a second messenger (Qi et al., 2022; Chen et al., 2025). These findings suggest that auxin receptors may function as multifunctional signaling hubs capable of integrating distinct regulatory inputs. In this context, NO-mediated redox regulation of TIR1 may represent an additional layer of control acting alongside other mechanisms that modulate receptor activity and stability. Such receptor multifunctionality expands the canonical view of auxin perception and raises intriguing questions about how plants coordinate redox signaling and cAMP-dependent pathways to achieve developmental plasticity during thermomorphogenesis.

At the physiological level, our findings provide direct evidence that NO participates in auxin-mediated hyponasty regulation (Figs. 2 and 5). Although hyponasty is a conserved adaptive response to diverse environmental cues, the contribution of NO has remained largely unresolved. In *A. thaliana*, hypoxia-induced NO accumulation has been linked to ethylene-dependent hyponastic movement (Hebelstrup et al., 2012). Our results suggest that NO may also contribute to hyponasty through TIR1-dependent auxin signaling, potentially linking redox regulation of auxin perception to plant responses across multiple environmental contexts. Interestingly, NO-mediated regulation of TIR1 has differential consequences in roots and shoots, uncovering an unexpected layer of tissue-specific regulation within the thermomorphogenic auxin signaling network. While the tir1C140A variant displays clear defects in hypocotyl elongation and hyponasty under warm temperatures, primary root growth appears largely unaffected under the same conditions (Fig. 5). Consistently, reducing endogenous NO levels or expressing the non-nitrosylatable tir1C140A variant only slightly affects primary root elongation and lateral root formation at 22 °C and has no detectable impact at 29 °C (Supplementary Fig. S7 and S8). These observations suggest that additional regulatory mechanisms may compensate for the absence of NO-dependent regulation of TIR1 in roots during thermomorphogenic responses.

One possible explanation for the differential requirement for TIR1 redox regulation between shoots and roots lies in the distinct cellular bases underlying growth in these organs. In Arabidopsis, hypocotyl thermomorphogenesis relies primarily on cell expansion, as cell division largely ceases after embryogenesis, whereas primary root growth depends on the combined contribution of cell elongation and sustained cell division within the root meristem. Although auxin regulates both processes (Perrot-Rechenmann, 2010), temperature reshapes auxin-responsive transcriptional programs differently in roots and shoots (Bianchinamo et al., 2023). Consistent with this organ-specific regulation, the transcription factor HY5 represses shoot thermomorphogenesis while promoting root growth under warm temperatures through the activation of auxin-responsive genes (Lee et al., 2021). Moreover, root and shoot thermomorphogenesis rely on partially overlapping but distinct temperature-signaling pathways: thermosensors that regulate shoot thermomorphogenesis do not appear to control primary root growth or branching, and roots can perceive temperature independently of the shoot (Bellstaedt et al., 2019; Borniego et al., 2022). Together, these differences suggest that root thermomorphogenesis is governed by additional regulatory layers controlling meristem activity, which may buffer the impact of modifications affecting TIR1 receptor function.

Divergence between root and shoot responses may also arise from differences in auxin receptor regulation and broader signaling networks. While S-nitrosylation of TIR1 and AFB2 has been linked to SCF complex assembly and Aux/IAA interaction, AFB1 appears to function through a distinct mechanism characterized by predominant cytosolic localization and reduced association with canonical SCF complexes (Chen et al., 2023; Dubey et al., 2023). Consistent with this view, NO influences root development under control temperatures, likely by promoting S-nitrosylation-dependent assembly of the SCF^TIR1–AFB2^ E3 ligase complex, whereas temperature has been shown to modulate the nucleocytoplasmic partitioning of AFB1 during primary root growth (Borniego et al., 2026). Hormonal crosstalk may further contribute to this divergence, as brassinosteroids regulate both meristem activity and cell elongation in roots and interact extensively with auxin signaling pathways (Nolan et al., 2020; Nolan et al., 2023).

Finally, the contribution of NO signaling is likely broader than the modification of a single protein target. S-nitrosylation regulates multiple components of hormone signaling networks, and individual proteins may harbor cysteine residues with distinct sensitivities to cellular NO levels (Paris et al., 2013). Such mechanisms have been described for regulatory proteins, including ASK1 and COP1, where S-nitrosylation at specific residues modulates E3 ligase activity in a context-dependent manner (Iglesias et al., 2018; Iglesias et al., 2024). Distinct NO dynamics and target repertoires in roots and shoots may therefore shape organ-specific thermomorphogenic responses by selectively modulating auxin receptor activity and associated signaling networks.

In summary, our work highlights how redox-dependent post-translational mechanisms may dynamically tune hormone perception to coordinate developmental responses to fluctuating environments. Understanding how NO signaling interfaces with auxin receptors and other regulatory networks may therefore provide new avenues for deciphering and potentially enhancing plant adaptive capacity in the context of climate warming. The evolutionary conservation of the Cys140 residue across plant species (Supplementary Fig. S5) further suggests that NO-dependent modulation of auxin perception may represent a broadly conserved mechanism contributing to growth plasticity. From a translational perspective, modulating TIR1 sensitivity to NO and cellular redox fluctuations could provide a strategic framework for fine-tuning auxin-dependent growth traits under elevated temperatures. Consequently, our findings offer a biochemical entry point for future studies aimed at enhancing thermotolerance and yield stability in crops facing increasingly warming climates.

## Acknowledgements

NCA, GM, DFF, MJI, and MCT are members of the research staff from CONICET. NMT is a doctoral fellow from the same institution.

## References

Bellstaedt, J., Trenner, J., Lippmann, R., Poeschl, Y., Zhang, X., Friml, J., Quint, M., Delker, C., 2019. A Mobile Auxin Signal Connects Temperature Sensing in Cotyledons with Growth Responses in Hypocotyls. Plant Physiol. 180, 757–766. 10.1104/pp.18.01377

Bianchimano, L., De Luca, M.B., Borniego, M.B., Iglesias, M.J., Casal, J.J., 2023. Temperature regulation of auxin-related gene expression and its implications for plant growth. J. Exp. Bot. 74, 7015–7033. 10.1093/jxb/erad265

Borniego, M.B., Costigliolo-Rojas, C., Casal, J.J., 2022. Shoot thermosensors do not fulfil the same function in the root. New Phytol. 236, 9–14. 10.1111/nph.18332

Borniego, M.B., Pereyra, M.E., Rossi, A., Sagerman-Furnas, K., Wilkinson, E., Strader, L., Casal, J.J., 2026. Thermosensory reconfiguration of the auxin transcriptional pathway to drive root cell growth. Nature Com, #NCOMMS-24-57040D, in press.

Cao, Z., Duan, X., Yao, P., Cui, W., Cheng, D., Zhang, J., Jin, Q., Chen, J., Dai, T., Shen, W., 2017. Hydrogen gas is involved in auxin-induced lateral root formation by modulating nitric oxide synthesis. International Journal of Molecular Sciences, 18(10), 2084. 10.3390/ijms18102084

Chen, H., Li, L., Zou, M., Qi, L., Friml, J. 2023. Distinct functions of TIR1 and AFB1 receptors in auxin signaling. Molecular Plant, 16(7), 1117–1119. doi: 10.1016/j.molp.2023.06.007.

Chen, H., Qi, L., Zou, M., Lu, M., Kwiatkowski, M., Pei, Y., Jaworski, K., Friml, J., 2025. TIR1-produced cAMP as a second messenger in transcriptional auxin signalling. Nature 640, 1011–1016. 10.1038/s41586-025-08669-w

Correa-Aragunde, N., Graziano, M., Lamattina, L., 2004. Nitric oxide plays a central role in determining lateral root development in tomato. Planta 218, 900–905. 10.1007/s00425-003-1172-7

Crawford, A.J., McLachlan, D.H., Hetherington, A.M., Franklin, K.A., 2012. High temperature exposure increases plant cooling capacity. Curr. Biol. 22, R396–R397. 10.1016/j.cub.2012.03.044

Dubey, S. M., Han, S., Stutzman, N., Prigge, M. J., Medvecká, E., Platre, M. P., Busch, W., Fendrychl, M., Estelle, M. 2023. The AFB1 auxin receptor controls the cytoplasmic auxin response pathway in Arabidopsis thaliana. Molecular Plant, 16(7), 1120–1130. 10.1016/j.molp.2023.06.008

Erwin, J.E., Heins, R.D., Karlsson, M.G., 1989. Thermomorphogenesis in Lilium Longiflorum. Am. J. Bot. 76, 47–52. 10.1002/j.1537-2197.1989.tb11283.x

Gaillochet, C., Burko, Y., Platre, M.P., Zhang, L., Simura, J., Willige, B., Kumar, S.V., Ljung, K., Chory, J., Busch, W., 2020. HY5 and phytochrome activity modulate shoot-to-root coordination during thermomorphogenesis in Arabidopsis. Development 147, dev192625. 10.1242/dev.192625

Gray, W.M., Del Pozo, J.C., Walker, L., Hobbie, L., Risseeuw, E., Banks, T., Crosby, W.L., Yang, M., Ma, H., Estelle, M., 1999. Identification of an SCF ubiquitin-ligase complex required for auxin response in *Arabidopsis thaliana*. Genes Develop. 13, 1678–1691. 10.1101/gad.13.13.1678

Gray, W.M., Östin, A., Sandberg, G., Romano, C.P., Estelle, M., 1998. High temperature promotes auxin-mediated hypocotyl elongation in Arabidopsis. Proc. Natl. Acad. Sci. U.S.A. 95, 7197–7202. 10.1073/pnas.95.12.7197

Hanzawa, T., Shibasaki, K., Numata, T., Kawamura, Y., Gaude, T., Rahman, A., 2013. Cellular Auxin Homeostasis under High Temperature Is Regulated through a SORTING NEXIN1-Dependent Endosomal Trafficking Pathway. Plant Cell 25, 3424–3433.

Hebelstrup, K.H., van Zanten, M., Mandon, J., Cristescu, S.M., Møller, I.M., Mur, L.A.J., 2012. Haemoglobin modulates NO emission and hyponasty under hypoxia-related stress in *Arabidopsis thaliana*. J. Exp. Bot., 63, 5581–5591. 10.1093/jxb/ers210

Huang, J., Wu, Q., Jing, H.K., Shen, R.F., Zhu, X.F., 2022. Auxin facilitates cell wall phosphorus reutilization in a nitric oxide-ethylene dependent manner in phosphorus deficient rice (*Oryza sativa L*.). Plant Sci. 322, 111371. 10.1016/j.plantsci.2022.111371

Iglesias, M.J., Costigliolo Rojas, C., Bianchimano, L., Legris, M., Schön, J., Gergoff Grozeff, G.E., Bartoli, C.G., Blázquez, M.A., Alabadí, D., Zurbriggen, M.D., Casal, J.J., 2024. Shade-induced ROS/NO reinforce COP1-mediated diffuse cell growth. Proc. Natl. Acad. Sci. U.S.A. 121, e2320187121. 10.1073/pnas.2320187121

Iglesias, M.J., Terrile, M.C., Correa-Aragunde, N., Colman, S.L., Izquierdo-Álvarez, A., Fiol, D.F., París, R., Sánchez-López, N., Marina, A., Calderón Villalobos, L.I.A., Estelle, M., Lamattina, L., Martínez-Ruiz, A., Casalongué, C.A., 2018. Regulation of SCFTIR1/AFBs E3 ligase assembly by S-nitrosylation of Arabidopsis SKP1-like1 impacts on auxin signaling. Redox Biol. 18, 200–210. 10.1016/j.redox.2018.07.003

Koini, M.A., Alvey, L., Allen, T., Tilley, C.A., Harberd, N.P., Whitelam, G.C., Franklin, K.A., 2009. High Temperature-Mediated Adaptations in Plant Architecture Require the bHLH Transcription Factor PIF4. Current Biol. 19, 408–413. 10.1016/j.cub.2009.01.046

Kolbert, Z., Bartha, B., Erdei, L., 2008. Exogenous auxin-induced NO synthesis is nitrate reductase-associated in *Arabidopsis thaliana* root primordia. Journal of plant physiology, 165(9), 967–975. 10.1016/j.jplph.2007.07.019

Kolbert, Z., Erdei, L., 2008. Involvement of nitrate reductase in auxin-induced NO synthesis. Plant Signaling & Behavior, 3(11), 972–973. 10.4161/psb.6170

Lee, S., Wang, W., Huq, E. 2021. Spatial regulation of thermomorphogenesis by HY5 and PIF4 in Arabidopsis. Nature communications, 12(1), 3656. 10.1038/s41467-021-24018-7

Lu, B., Wang, S., Feng, H., Wang, J., Zhang, K., Li, Y., Wu, P., Zhang, M., Xia, Y., Peng, C., Li, C., 2024. FERONIA-mediated TIR1/AFB2 oxidation stimulates auxin signaling in Arabidopsis. Mol. Plant 17, 772–787. 10.1016/j.molp.2024.04.002

Manafi, H., Baninasab, B., Gholami, M., Talebi, M., 2021. Nitric oxide induced thermotolerance in strawberry plants by activation of antioxidant systems and transcriptional regulation of heat shock proteins. The Journal of Horticultural Science and Biotechnology, 96(6), 783–796. 10.1080/14620316.2021.1927206

Muñoz, A., Mangano, S., Toribio, R., Fernández-Calvino, L., Del Pozo, J.C., Castellano, M.M., 2022. The co-chaperone HOP participates in TIR1 stabilization and in auxin response in plants. Plant Cell Environ. 45, 2508–2519. 10.1111/pce.14366

Nolan, T. M., Vukašinović, N., Liu, D., Russinova, E., Yin, Y. 2020. Brassinosteroids: multidimensional regulators of plant growth, development, and stress responses. The plant cell, 32(2), 295–318. 10.1105/tpc.19.00335

Nolan, T. M., Vukašinović, N., Hsu, C. W., Zhang, J., Vanhoutte, I., Shahan, R., Taylor, I.W., Greenstreet, L., Heitz, M., Afanassiev, A., Wang, P., Szekely, P., Brosnan, A., Yin, Y., Schiebinger, G., Russinova, E., Benfey, P. N. 2023. Brassinosteroid gene regulatory networks at cellular resolution in the Arabidopsis root. Science, 379(6639), eadf4721. 10.1126/science.adf4721

París, R., Iglesias, M. J., Terrile, M. C., Casalongué, C. A. 2013. Functions of S-nitrosylation in plant hormone networks. Frontiers in plant science, 4, 294.10.3389/fpls.2013.00294

París, R., Vazquez, M.M., Graziano, M., Terrile, M.C., Miller, N.D., Spalding, E.P., Otegui, M.S., Casalongué, C.A., 2018. Distribution of Endogenous NO Regulates Early Gravitropic Response and PIN2 Localization in Arabidopsis Roots. Front. Plant Sci. 9, 495. 10.3389/fpls.2018.00495

Perrot-Rechenmann, C., 2010. Cellular Responses to Auxin: Division versus Expansion. Cold Spring Harbor Perspectives in Biology 2, a001446–a001446. 10.1101/cshperspect.a001446

Piterková, J., Luhová, L., Mieslerová, B., Lebeda, A., Petřivalský, M., 2013. Nitric oxide and reactive oxygen species regulate the accumulation of heat shock proteins in tomato leaves in response to heat shock and pathogen infection. Plant Science, 207, 57–65. 10.1016/j.plantsci.2013.02.010

Pucciariello, O., Legris, M., Costigliolo Rojas, C., Iglesias, M.J., Hernando, C.E., Dezar, C., Vazquez, M., Yanovsky, M.J., Finlayson, S.A., Prat, S., Casal, J.J., 2018. Rewiring of auxin signaling under persistent shade. Proc. Natl. Acad. Sci. U.S.A. 115, 5612–5617. 10.1073/pnas.1721110115

Qi, L., Kwiatkowski, M., Chen, H., Hoermayer, L., Sinclair, S., Zou, M., del Genio, C.I., Kubeš, M.F., Napier, R., Jaworski, K., Friml, J., 2022. Adenylate cyclase activity of TIR1/AFB auxin receptors in plants. Nature 611, 133–138. 10.1038/s41586-022-05369-7

Sánchez-Vicente, I., Lorenzo, O., 2021. Nitric oxide regulation of temperature acclimation: a molecular genetic perspective. Journal of Experimental Botany, 72(16), 5789–5794. 10.1093/jxb/erab049

Tao, Y., Ferrer, J.-L., Ljung, K., Pojer, F., Hong, F., Long, J.A., Li, L., Moreno, J.E., Bowman, M.E., Ivans, L.J., Cheng, Y., Lim, J., Zhao, Y., Ballaré, C.L., Sandberg, G., Noel, J.P., Chory, J., 2008. Rapid Synthesis of Auxin via a New Tryptophan-Dependent Pathway Is Required for Shade Avoidance in Plants. Cell 133, 164–176. 10.1016/j.cell.2008.01.049

Tatematsu, K., Kumagai, S., Muto, H., Sato, A., Watahiki, M.K., Harper, R.M., Liscum, E., Yamamoto, K.T., 2004. MASSUGU2 Encodes Aux/IAA19, an Auxin-Regulated Protein That Functions Together with the Transcriptional Activator NPH4/ARF7 to Regulate Differential Growth Responses of Hypocotyl and Formation of Lateral Roots in Arabidopsis thaliana. Plant Cell 16, 379–393. 10.1105/tpc.018630

Terrile, M.C., París, R., Calderón-Villalobos, L.I.A., Iglesias, M.J., Lamattina, L., Estelle, M., Casalongué, C.A., 2012. Nitric oxide influences auxin signaling through S-nitrosylation of the Arabidopsis TRANSPORT INHIBITOR RESPONSE 1 auxin receptor. Plant J. 70, 492–500. 10.1111/j.1365-313X.2011.04885.x

Terrile, M.C., Tebez, N.M., Colman, S.L., Mateos, J.L., Morato-López, E., Sánchez-López, N., Izquierdo-Álvarez, A., Marina, A., Calderón Villalobos, L.I.A., Estelle, M., Martínez-Ruiz, A., Fiol, D.F., Casalongué, C.A., Iglesias, M.J., 2022. S-Nitrosation of E3 Ubiquitin Ligase Complex Components Regulates Hormonal Signalings in Arabidopsis. Frontiers Plant Sci. 12.

Ulmasov, T., Murfett, J., Hagen, G., Guilfoyle, T.J., 1997. Aux/IAA proteins repress expression of reporter genes containing natural and highly active synthetic auxin response elements. Plant Cell 9, 1963–1971. 10.1105/tpc.9.11.1963

Van Zanten, M., Voesenek, L.A.C.J., Peeters, A.J.M., Millenaar, F.F., 2009. Hormone- and Light-Mediated Regulation of Heat-Induced Differential Petiole Growth in Arabidopsis. Plant Physiol. 151, 1446–1458. 10.1104/pp.109.144386

Wang, R., Zhang, Y., Kieffer, M., Yu, H., Kepinski, S., Estelle, M., 2016. HSP90 regulates temperature-dependent seedling growth in Arabidopsis by stabilizing the auxin co-receptor F-box protein TIR1. Nat Commun 7, 10269. 10.1038/ncomms10269

Watanabe, E., Mano, S., Nomoto, M., Tada, Y., Hara-Nishimura, I., Nishimura, M., Yamada, K., 2016. HSP90 Stabilizes Auxin-Responsive Phenotypes by Masking a Mutation in the Auxin Receptor TIR1. Plant Cell Physiol. 57, 2245–2254. 10.1093/pcp/pcw170

Wilkinson, J.Q., Crawford, N.M., 1991. Identification of the Arabidopsis CHL3 gene as the nitrate reductase structural gene NIA2. Plant Cell 3, 461–471. 10.1105/tpc.3.5.461

Yadav, S., David, A., Bhatla, S.C., 2011. Nitric oxide accumulation and actin distribution during auxin-induced adventitious root development in sunflower. Sci. Hortic.129, 159–166. 10.1016/j.scienta.2011.03.030

Yang, X., Dong, G., Palaniappan, K., Mi, G., Baskin, T.I., 2017. Temperature-compensated cell production rate and elongation zone length in the root of Arabidopsis thaliana. Plant Cell Environ. 40, 264–276. 10.1111/pce.12855

Yu, H., Zhang, Y., Moss, B.L., Bargmann, B.O.R., Wang, R., Prigge, M., Nemhauser, J.L., Estelle, M., 2015. Untethering the TIR1 auxin receptor from the SCF complex increases its stability and inhibits auxin response. Nat. Plants 1, 14030. 10.1038/nplants.2014.30

Zhang, X.-C., Millet, Y.A., Cheng, Z., Bush, J., Ausubel, F.M., 2015. Jasmonate signalling in Arabidopsis involves SGT1b–HSP70–HSP90 chaperone complexes. Nat. Plants 1, 15049. 10.1038/nplants.2015.49

