## Supplemental figures for "Nitric Oxide Modulates Auxin Signaling through TIR1 S-Nitrosylation During Thermomorphogenesis in Arabidopsis"

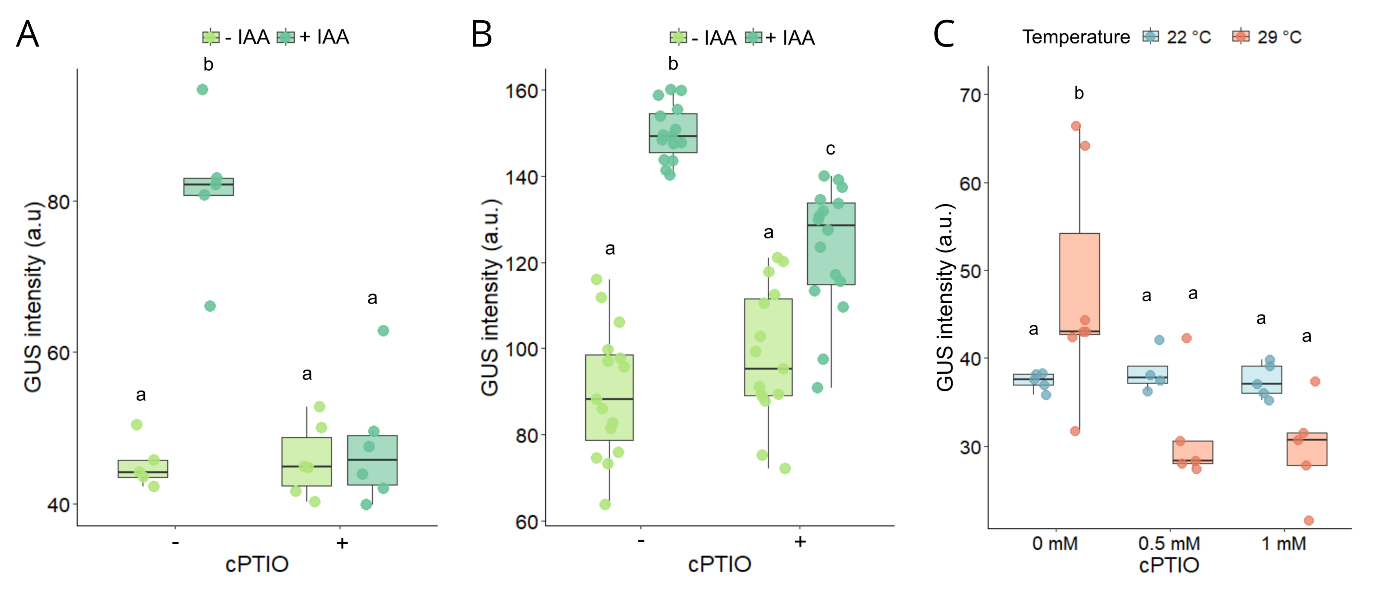

**Figure S1. Quantification of GUS signal intensity in response to IAA, temperature, and cPTIO.** GUS activity is expressed in arbitrary units (a.u.). Boxplots show the median and quartiles, with individual dots representing independent biological replicates. **(A)** Hypocotyls of 3-day-old IAA19:GUS seedlings treated with 10 µM IAA and/or 1 mM cPTIO for 2 h. (B) Root tips of 4-day-old DR5:GUS seedlings treated with 1 µM IAA and/or 1 mM cPTIO for 2 h. (C) Hypocotyls of 3-day-old IAA19:GUS seedlings grown at 22 °C or 29 °C and treated with 0, 0.5, or 1 mM cPTIO. Statistical significance was determined by two-way ANOVA followed by Tukey’s HSD post-hoc test (p < 0.05); different letters indicate significant differences between treatments.

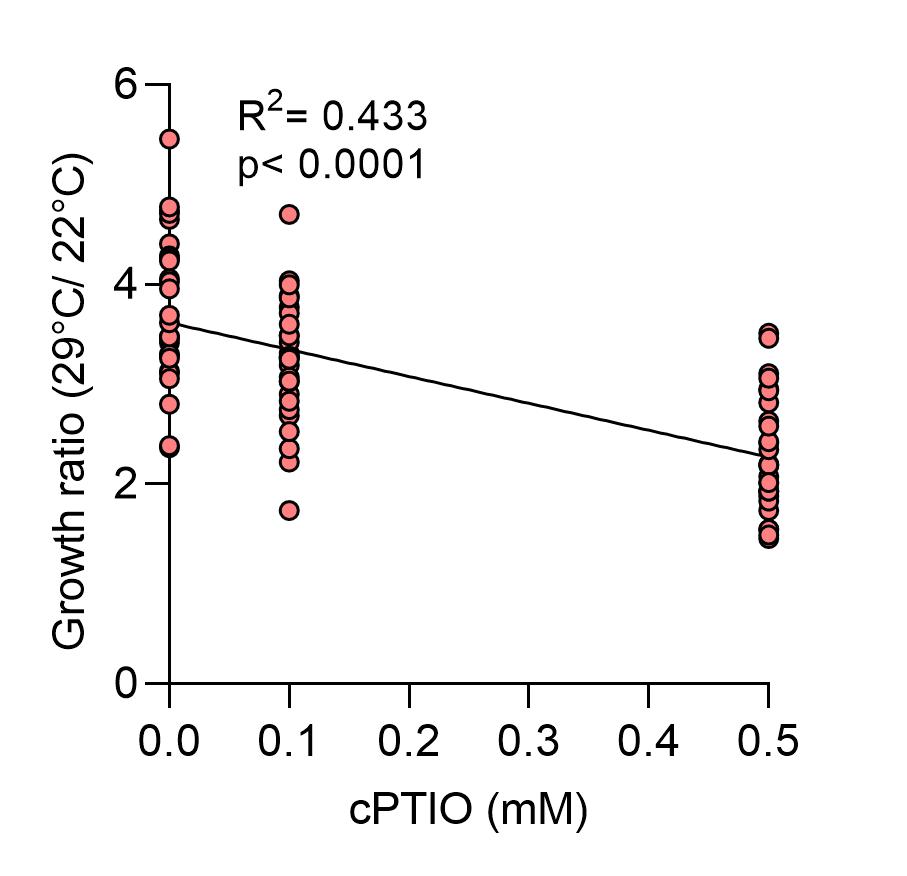

**Figure S2: Hypocotyl elongation is inhibited by cPTIO in a dose-dependent manner under warmth.** Linear regression adjustment of growth rate of Arabidopsis hypocotyls to increasing concentrations of cPTIO under warmth. The significance of the R2 is indicated.

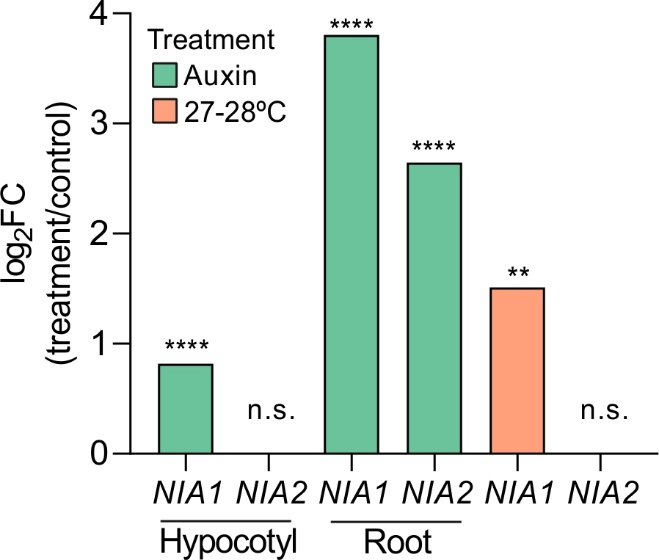

**Figure S3. Expression of *Nitrate Reductase (NR)* genes in response to auxin and temperature.** Log fold changes (log_2_FC) of *NIA1* (AT1G77760) and *NIA2* (AT1G37130) expression were compiled from publicly available transcriptomic datasets. Auxin-responsive expression in hypocotyls and roots of 5 day-old seedlings was obtained from microarray and RNA-seq experiments (Chapman et al., 2013, PloS one, 7(5), e36210; Omelyanchuk et al., 2017, Scientific reports, 7(1), 2489); while temperature-responsive expression in shoots was derived from a published meta-analysis (Bianchimano et al 2023). In each case, log2FC values were calculated relative to the corresponding control condition reported in the original studies. Significant differential expressions (as defined in the original analyses) are shown. *** p<0.001 ****p<0.0001.

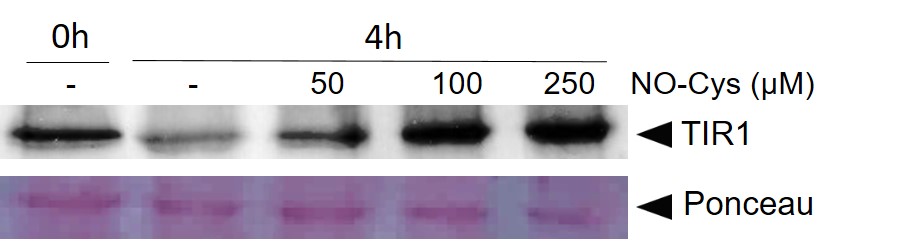

**Figure S4: TIR1 degradation is inhibited by NO donor in a dose-response manner.** *In vitro* degradation assays were performed using protein extracts from *tir1-1* 35S:TIR1-myc five-day-old seedlings incubated for 4 h in the presence of increasing concentrations of NO donor NO-Cys. Protein extracts were analyzed by Western blot, and TIR1-VENUS levels were quantified relative to RUBISCO (Ponceau staining) used as a loading control.

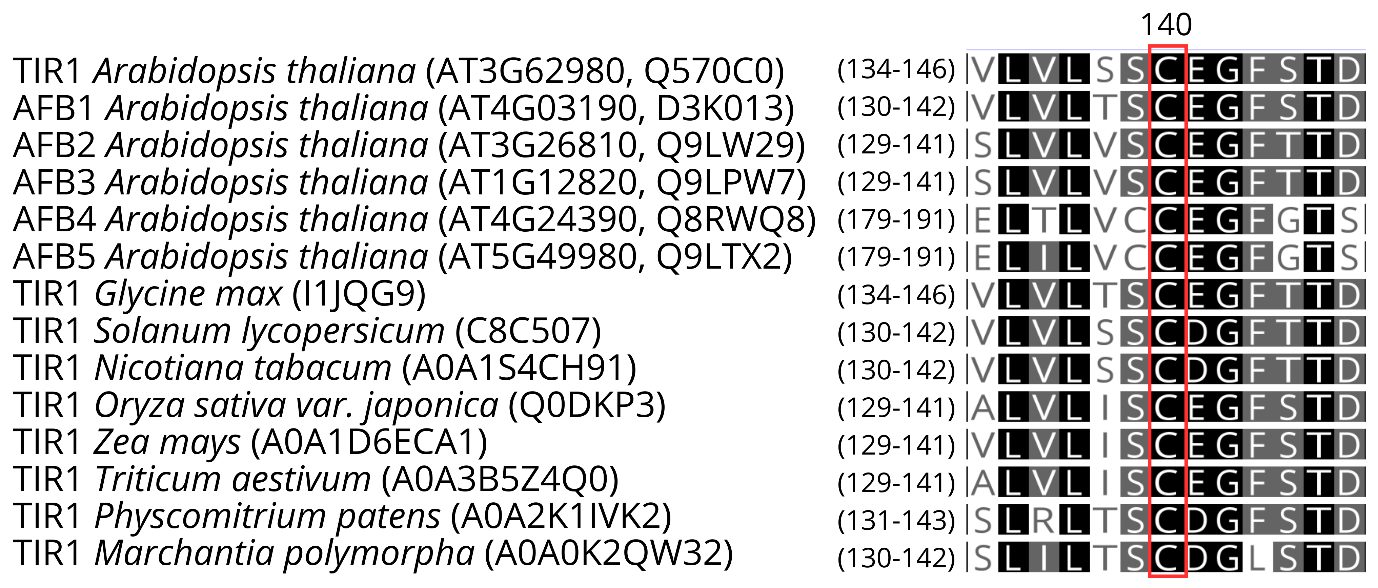

**Figure S5. Evolutionary conservation of the Cys140 residue in TIR1/AFB homologs across diverse plant species.** Multiple sequence alignment of amino acid sequences from Arabidopsis thaliana auxin receptors (TIR1 and AFB1-5) and TIR1 orthologs from agronomically important species and bryophytes. For each sequence, the corresponding TAIR or UniProt accession numbers are provided. The numbers in parentheses preceding the alignment indicate the specific amino acid positions represented for each protein. The red box highlights the absolute conservation of the cysteine residue corresponding to position 140 in AtTIR1.Protein sequences were retrieved from the NCBI, TAIR, and UniProt databases and aligned using Geneious software. Shading indicates amino acid similarity levels (Black: 100% similarity; Gray: 80% to 100% similarity; White: less than 60% similarity).

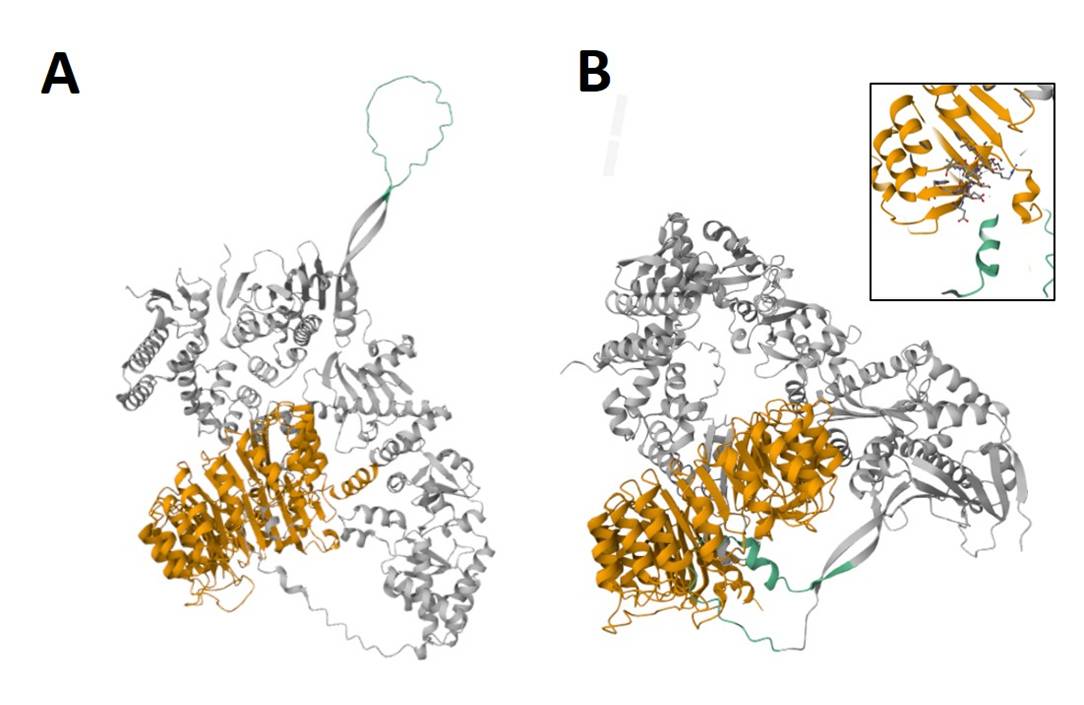

**Figure S6: Effect of TIR1 S-nitrosylation on the TIR1–HSP90 interaction.** Cartoon representations of the best structural models predicted by AlphaFold3 for the interaction between SKP1, HSP90, and TIR1 **(A)** or S-nitrosylated TIR1 at Cys140 (Cys140-SNO-TIR1) **(B)**. The S-nitrosylated cysteine residue (Cys140-SNO; shown in blue, atomic representation) is highlighted in the inset of panel B. Confidence scores for the predicted interface. The inclusion of the S-nitrosothiol group at Cys140 increased the interface Predicted Template Modeling (ipTM) score from 0.48 to 0.53. TIR1 is shown in orange, and the HSP90 segment (residues 230–250) is shown in green. SKP1 and the remaining portion of the HSP90 structure are shown in grey. Models correspond to the top-ranked AlphaFold3 predictions obtained using the multimer modeling mode. Models correspond to the top-ranked AlphaFold3 predictions obtained using the multimer modeling mode.

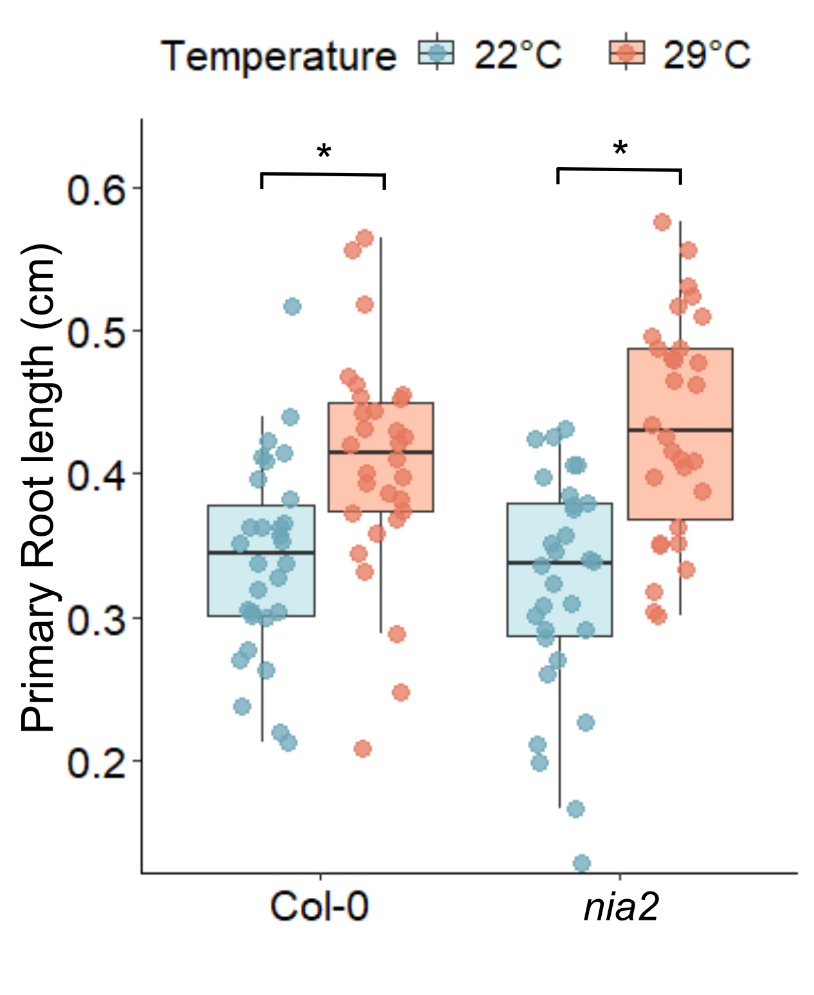

**Figure S7: Primary root response to warmth in *nia2* mutant.** Four-day-old Col-0 and *nia2* seedlings were transferred to 22°C or 29°C, and primary root length was measured after 2 days. Boxplots represent the median and the first and third quartiles, and the whiskers extend to minimum and maximum values; all data points are shown as dots. Statistical differences were assessed using a generalized linear model (GLM, gamma distribution) followed by Tukey’s multiple comparison test (p < 0.05), as indicated by letters.

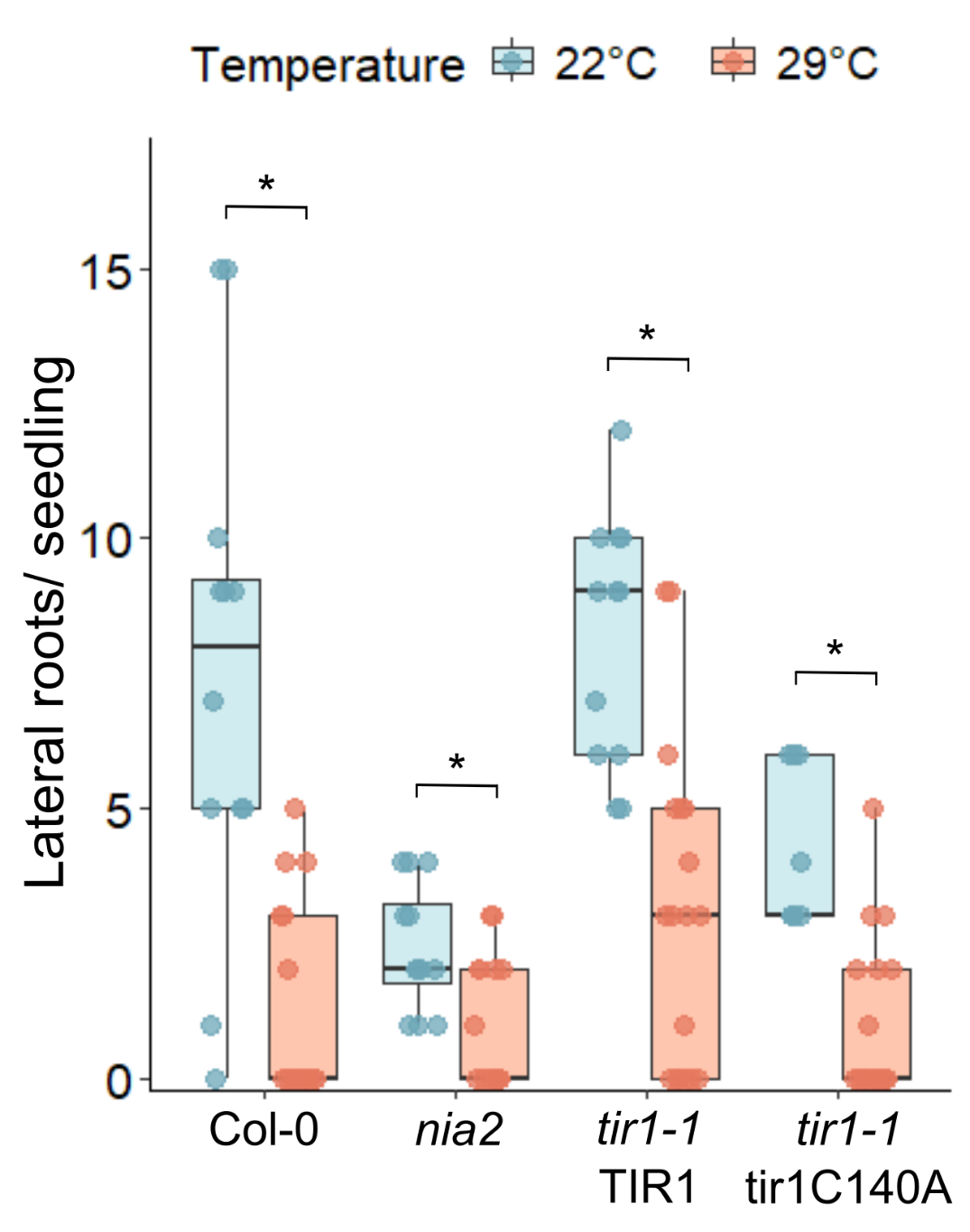

**Figure S8: LR development under warmth is not regulated by NO.** Four-day-old Col-0, *nia2, tir1-1, tir1-1* TIR1-myc *and tir1-1* tir1C140A-my*c* seedlings were transferred to 22°C or 29°C, and primary root length was measured after 5 days. Boxplots represent the median and the first and third quartiles, and the whiskers extend to minimum and maximum values; all data points are shown as dots. Statistical differences were assessed using a generalized linear model (GLM, gamma distribution) followed by Tukey’s multiple comparison test (p < 0.05), as indicated by letters.
